# Inferring Gene Presence in Incomplete Data via Phylogenetic Occupancy Modeling

**DOI:** 10.64898/2026.02.27.708499

**Authors:** John S.A. Mattick, Wesley C. DeMontigny, Charles F. Delwiche

**Author notes:** Equally contributing authors (John S.A. Mattick, Wesley C. DeMontigny).

## Abstract

Increasing access to genomic data has revolutionized our understanding of biology. Organisms that were previously unculturable or otherwise difficult to study have been investigated using metagenomic sequencing and bioinformatic assemblies, illuminating biological diversity that was previously invisible. However, as the availability of genomic data has grown, so has the challenge posed by incomplete genomes. Many genomes obtained from metagenomic assemblies or mixed cultures are of poor quality and establishing fully complete genomes requires substantial effort. Incomplete genomes pose difficulties for several common analyses, particularly gene-inventory and core-genome analyses. When genomes are incomplete, distinguishing true gene absence from non-detection becomes difficult. For relatively complete genomes, gene absences are inferred to be true absences, but for highly incomplete genomes, researchers often exclude such data entirely. Probabilistic models have attempted to address this issue in core genome analyses. In particular, mOTUpan utilizes an iterative algorithm to infer genome completeness and categorize genes as core or accessory. In this work, we substantially improve upon this approach by integrating a well-established class of ecological models, called occupancy models, with evolutionary modeling. Our “phylogenetic occupancy model” defines a probability distribution over gene presence that accounts for shared information across related genomes. This framework simultaneously estimates genome completeness and the probability that a gene is present but unobserved. This model substantially outperforms competing methods for core genome inference and enables inference of single-gene presence/absence and ancestral-state reconstruction. Alongside this paper, we provide our model as a Python package.

## Introduction

Our understanding of the microbial world has been revolutionized several times, most recently by the metagenomics-driven expansion of the catalog of Earth’s diversity (1). The advent of the microscope enabled the discovery and morphological characterization of microorganisms, which were eventually classified into two major categories: eukaryotes and prokaryotes (2). Later, the discovery of DNA and the advent of sequencing technologies enabled the characterization of the genomic content of these organisms and the use of those genomes to define relationships based on shared genetic history, in addition to shared morphology (3). Small subunit ribosomal RNA (16S rRNA), which has been used for over 50 years to define the relationships between different species of bacteria (4, 5), was able to identify a third domain of life, the Archaea, whose morphology superficially resembles bacteria, but that are genetically closer to eukaryotes (6, 7). As sequencing costs have decreased, genomics has shifted from the characterization of isolated single organisms to broader sequencing projects that study entire microbial communities. Metagenomics is the study of genetic material directly recovered from environmental samples and has redefined our understanding of microbial diversity on Earth (8–10). The number of cultured (or culturable) microorganisms remains unclear, but some studies estimate that as few as 0.5% of bacteria and archaea can be cultured using standard techniques (11). Metagenomic analysis has led to the discovery of thousands of novel taxa and clades, and revolutionized our understanding of the tree of life.

One of the most powerful tools in metagenomics is the ability to assess the genomic content of hundreds of genomes from a specific taxon without isolating and growing them individually (12). However, the genomes acquired by these methods are often incomplete, either due to low sequencing depth or difficulty in taxonomic binning (13); even deep sequencing can yield partial genomes of rare taxa. In fact, this is representative of a broader trend in genomics, in which the number of partial genomic assemblies vastly outnumbers that of complete or near-complete assemblies (14).

To understand the functional capabilities of a given clade, gene inventories and core genome analyses were developed. Gene inventories examine the presence or absence of specific gene families to characterize the functional capacities of individual taxa, whereas core genome analyses identify the set of genes shared by all members of a clade to infer their common biological capabilities (15). However, incomplete genomic data make these analyses challenging, since the absence of a gene from a genome assembly does not necessarily imply its absence from the true genome (16). Explicit treatment of incomplete information in gene inventories remains uncommon, and core genome analyses have largely relied on inclusion thresholds or empirical gene presence thresholding to accommodate variability in genome completeness. Other researchers have attempted to leverage the shared information between genomes to make more informed inferences about the core genomes in the presence of incomplete data. PPanG-GOLiN (17), for example, uses synteny networks to identify clusters of co-occurring genes and classifies genes into “persistent,” “shell,” or “cloud” populations. Although this approach is highly scalable, its reliance on synteny is problematic for genomes that are highly fragmented or so distantly related that synteny is no longer conserved. mOTUpan (18) adopts a probabilistic approach to core genome inference, using an iterative algorithm that classifies genes as either core or accessory. Core gene presence is modeled as a function of genome completeness, whereas accessory genes are assumed to be independently distributed across genomes, following a gene-frequency-dependent distribution informed by inferred genome sizes. While this performs well at the low evolutionary distances for which it was designed, the assumption that core genes are those whose presence-absence patterns are explained entirely by genome completeness becomes problematic at deeper evolutionary depths. Moreover, if the dataset is heavily biased toward particular sub-clades, this approach is likely to produce a substantial number of false-positive core genes.

The model used by mOTUpan can be viewed as an occupancy model. These models are popular in ecology for assessing the probability that a species occupies a particular site given incomplete (or non-existent) survey data at that site, given repeated observations (19–22). In general, occupancy models assign a probability that species *i* occupies site *j* given an observation *x*_*ij*_ as:

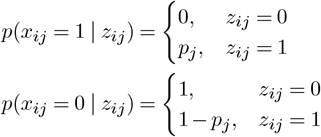

where *p*_*j*_ indicates the completeness of the survey at site *j* and *z*_*ij*_ is the true occupancy state, with *z*_*ij*_ = 1 if the species truly occupies that site and *z*_*ij*_ = 0 otherwise. In the case of mOTUpan, there is a mixture of models: one component models genes as being distributed according to their frequency of occurrence (the accessory component), and another component acts as an occupancy model, where the true state *z*_*ij*_ is 1 for all genes in that mixture component. There are several areas where this model could be improved. First, even genes that exhibit seemingly random distributions may contain valuable information about the completeness of a given genome. Second, not all genes whose presence is approximately predictable under an occupancy model belong in the “core” set. Indeed, the core set in the context of mOTU-pan is simply the set of genes whose presence is predictable by genome completeness, rather than the set of genes that are universal across all genomes.

Ideally, we would like to pool information across all genomes *j* according to their similarity. In ecological occupancy modeling, this can be accomplished using Gaussian processes to leverage spatial covariance among survey sites (23) or ecological covariates of the sites. In the context of genomes, however, these approaches are not particularly useful. There are no especially meaningful notions of distance between genomes, and designing covariates for each metagenomic dataset would require complex dataset-specific choices. An alternative approach is to use a belief network. Belief networks define complex joint distributions over random variables through a set of dependence relationships that form a directed acyclic graph (24). Such approaches have been used to model between-species dependence in ecological occupancy models (25), but are less commonly applied to survey sites. In the context of genomes, a natural choice for the belief network structure is a phylogenetic tree. Although statistical phylogenetic models are typically formulated as stochastic processes evolving along the branches of a tree, they may equivalently be viewed as belief networks. In this representation, two genomes are conditionally independent given the state of their most recent common ancestor, and the joint distribution over ancestral and descendant states is governed by a global set of shared parameters (i.e., the substitution rate parameters) together with relationship-specific parameters (i.e., the branch lengths) that control the divergence of the joint distribution from some limiting stationary distribution.

Designing our occupancy model as a belief network allows us to draw on a broad body of well-established literature and algorithms from probabilistic graphical modeling. In particular, it provides several options for the model’s output. In our case, both the sum-product and max-product algorithms seem particularly useful. These algorithms compute the marginal posterior probability that a gene is in a genome and the joint maximum *a posteriori* state across all genomes, respectively (26, 27). Because these algorithms propagate across the entire phylogenetic tree, they also identify the marginal probability and the maximum *a posteriori* state for each ancestral node. This allows us to better understand genomic content in extinct organisms when the data from their descendants is incomplete.

Therefore, to improve probabilistic assessments of gene presence-absence in incomplete genomic data, we developed a probabilistic model in which a phylogenetic belief network defines the joint distribution of the true presence-absence states across genomes, while a full occupancy model specifies the observation process. We benchmark this model for internal coherence using simulations, and evaluate its performance on empirical datasets from *α*-proteobacteria and *γ*-proteobacteria. We also showcase the ability of the model to reconstruct ancestral genome content on the phylogeny in the Asgardarchaea. Our approach outperforms comparable methods, such as mOTUpan and simple completeness thresholding, while enabling richer downstream analyses. In particular, the model assigns posterior probabilities to the presence of every gene in every genome (rather than merely assigning genes to categories) and naturally supports ancestral state reconstruction within the same probabilistic framework.

## Methods

### Phylogenetic Occupancy Modeling

In a phylogenetic occupancy model, our goal is to infer the true occupancy state *z*_*ij*_ for each gene *i* in each genome *j*. Each genome is associated with a completeness parameter *p*_*j*_, which governs the conditional distribution of the observed presence–absence data *x*_*ij*_ given the latent occupancy state. As described above, these conditional probabilities follow the standard occupancy model,

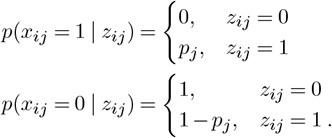

This construction already defines a simple belief network, but with dependencies restricted to pairs of variables (*z*_*ij*_, *x*_*ij*_). Our aim is to extend this structure so that, for a fixed gene *i*, the latent occupancy states (*z*_*i*1_, …, *z*_*iN*_) across genomes are coupled through a joint distribution that reflects their evolutionary relatedness, with more closely related genomes being more likely to share the same state.

We encode this dependence through a set of conditional independence relationships defined on a phylogenetic tree. For any two genomes *a* and *b*, the corresponding occupancy states for gene *i* are conditionally independent given the state at their most recent common ancestor. Conditional probabilities of occupancy state along each branch of the tree are modeled using a symmetric two-state process, with the conditional probabilities

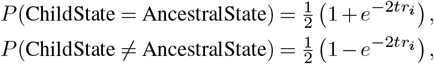

where *t* is a branch-specific parameter measuring the evolutionary distance between a node and its ancestor, traditionally interpreted as the elapsed time of a continuous-time Markov chain; this parameterization is chosen to ensure increasing agnosticism about the occupancy state as evolutionary distance grows, with the descendant state approaching a uniform distribution for large *t*. To account for the fact that gene families exhibit different levels of transience in a genome, we use a mixture of rate multipliers for *t*. In each mixture component, we assign a new parameter *r*_*i*_ that acts as a multiplicative factor for the branch lengths *t*. In this case, we set the values of *r*_*i*_ to come from a discretized standard log-normal distribution. This approach can be viewed as a modification of the phylogenetic method for Gamma site-rate heterogeneity (28). We found that simultaneously estimating the parameters of the discretized mixture distribution led to unstable optimization in NumPyro, whereas the standard log-normal mixture performed well. In all of our analyses, we utilize a 10-component mixture.

Likelihoods are computed by propagating these conditional probabilities down the phylogenetic tree using Felsenstein’s pruning algorithm (29). This computation is equivalent to exact marginalization via variable elimination on a tree-structured belief network or the sum-product algorithm, where the query node is only the root (26, 27, 30). At the root of the tree, instead of weighting by the stationary distribution of a Markov chain, we introduce a prior vector ***π*** that represents the proportion of genes present at the ancestral genome of the entire dataset. Thus, depending on the rooting of the tree, we are likely to infer a slightly different *π*, since it represents a prior probability assigned to a different node.

The model was implemented in Python using NumPyro (31, 32), enabling efficient maximum likelihood estimation via automatic differentiation and the ADAM optimizer. Preliminary experiments with Hamiltonian Monte Carlo in NumPyro indicated that the posterior distribution was tightly concentrated around the maximum likelihood estimate, but was much more computationally intensive. Given the large number of benchmarking runs performed here, we therefore report maximum likelihood estimates. We simultaneously estimate genome completeness ***p***, branch lengths ***t***, and ancestral proportion ***π*** = [*π*_0_, 1 − *π*_0_] given a dataset and a fixed phylogeny. Initial experimentation indicated that 1,000 epochs of optimization yielded a good fit on the datasets we investigated. Following parameter estimation, we apply the sum-product algorithm to compute marginal posterior probabilities of occupancy for each gene at each node and genome, and the max-product algorithm to obtain a joint maximum *a posteriori* reconstruction of occupancy states across all genomes. On a single NVIDIA L4 GPU, inference for a single dataset takes minutes. This makes data curation the most time-consuming part of these analyses.

The phylogenetic occupancy model can be interpreted as a statistical phylogenetic model with taxon-specific state ambiguity (e.g., utilizing IUPAC ambiguity codes in an amino-acid phylogeny). In this view, an observed *x*_*ij*_ = 1 corresponds to an unambiguous presence state, whereas *x*_*ij*_ = 0 is ambiguous, representing either true absence or unobserved presence, with the degree of ambiguity determined by the genome-specific occupancy model. In terms of a generative probabilistic model, this encodes a process whereby some proportion of genes 1 − *π*_0_ are present at the root of the phylogeny and are gained or lost across the tree according to a continuous-time Markov chain to produce a final true gene occupancy state *z*_*ij*_ for each gene *i* and each genome *j*. Finally, if the true occupancy state is *z*_*ij*_ = 0, we will observe *x*_*ij*_ = 0, however if *z*_*ij*_ = 1, we will observe *x*_*ij*_ = 1 with probability *p*_*j*_ and *x*_*ij*_ = 0 with probability 1 − *p*_*j*_.

### Simulation Analysis

Datasets with varying numbers of genomes and degrees of completeness were simulated under the model to characterize its behavior when the model was well specified. Specifically, datasets with 10, 50, 100, 500, and 750 genomes were simulated, each containing 5,000 potentially acquirable genes, with genome completeness values drawn from a Beta distribution. We considered 8 different parameterizations of the Beta distribution, spanning a wide range of completeness profiles, using the following treatments: (*α, β*) ∈ {(30, 10), (5, 1), (3, 0.5), (1, 1), (0.25, 0.25), (1, 5), (0.5, 3.0)}. For each combination of completeness profile and number of genomes, we simulated 10 datasets, each with a randomly generated phylogenetic tree, yielding a total of 400 analyses. The tree diameter of each phylogeny was set to be 0.5 for all simulations, meaning each ortholog has been gained/lost approximately 0.5 times from the root to the longest tip of the tree. For each simulation, model outputs were recorded, and precision and recall were computed under the different conditions. Here, we do not consider observed presence (*x*_*ij*_ = 1) in our precision/recall calculations, as these inferences are guaranteed to be correct. The results were visualized using ggplot2.

### Empirical Evaluations

*α*-proteobacterial and *γ*-proteobacterial genomes (442 and 694 genomes, respectively) were pulled from the NCBI genome database by selecting all genomes that were categorized under these taxonomic designations in NCBI and had the following qualities: reference genomes, annotated by NCBI RefSeq, from type materials, excluding atypical genomes, excluding metagenomic assemblies, and excluding genomes from large multi-isolate projects. Proteins from these genomes were assigned to orthogroups via the emapper tool in the EggNog2 suite (33, 34). Concatenated amino acid phylogenies were generated by identifying orthogroups that were present in 95% of genomes with a copy number of 1 or 2. When 2 copies were present, the longest ortholog was selected. Each orthogroup was aligned with MAFFT v.7.5.20 (35) using the –auto parameter and trimmed with TrimAl using the - automated1 parameter (36). These trimmed alignments were concatenated, and phylogenies were inferred with IQTree2 v. 2.2.7 (37), using models selected by IQTree2’s model finder. These phylogenies and high-quality gene presence/absence data were treated as ground truth to evaluate our model’s performance on biological data.

We artificially depleted orthogroups from our genomes using the same completeness profiles (Beta distributions) described above and analyzed the resulting datasets under our model. For each genome *j*, we draw a completeness parameter *p*_*j*_ and mask (1 − *p*_*j*_)% of the present orthogroups from the data. After each dataset was depleted, we discarded orthogroups present in fewer than 5 depleted genomes to simulate the removal of extremely rare orthogroups. We performed this procedure 10 times per completeness profile, yielding 80 analyses per dataset.

Using both the marginal probabilities of occupancy and the maximum *a posteriori* estimates, we inferred core genomes for major *α*-proteobacterial and *γ*-proteobacterial clades and assessed their performance against the observed ground truth. We compared the precision and recall of our core genome estimates with those of empirical presence-thresholding methods (i.e., genes present in 90% or 95% of observed incomplete genomes) and with mOTUpan predictions. We investigated both definitions of a true core genome, in which genes are present in all members of a clade, and a relaxed core genome, in which genes are present in at least 90% of members of a clade. The results were visualized using ggplot2.

### Asgardarchaea Analysis

The Asgardarchaeal dataset was generated from all Asgard genomes present in Genbank as of October 2025. Unlike the proteobacterial genomes, there was no discrimination in genome quality: all genomes were downloaded, including 304 with predicted protein sequences and 437 with only nucleic acid sequences. Genomes without proteins had their protein content inferred using Prokka with the –kingdom Archaea parameter (38). In addition, 206 reference-quality and complete genomes from the TACK group of Archaea were downloaded. A phylogeny based on a concatenated amino-acid alignment of the combined TACK and Asgard dataset was inferred, as was done for the *α*-proteobacterial and *γ*-proteobacterial datasets. Two Asgard genomes were excluded from the analysis because 90% of the orthogroups used to generate the phylogeny were missing from these genomes. The TACK outgroup was excluded from downstream analyses and included solely to infer a root for ancestral reconstruction within the Asgard clade. We analyzed this dataset in our phylogenetic occupancy model with particular focus on the ancestral states of the major lineages of the Asgards. We specifically focused on orthogroups identified as belonging to the set of eukaryotic-specific proteins (ESPs) identified by Liu et al (39). Proteins from the Asgard dataset were assigned to AsCoG designations and filtered to only include AsCogs identified as ESPs. These proteins then filtered the orthogroup list, retaining only orthogroups containing ESP-designated AsCog proteins for analysis in the ancestral state reconstruction. Joint reconstruction was used to infer ancestral genome content at each node on the phylogeny. Gains and losses along the tree were plotted using ggtrees for visualization (40). Specific nodes were selected as roots of major Asgard clades using NCBI species designations. The operational clade definitions for Heimdal, Loki, Hel, Thor, Odin, Hod, Njord, Wukong, Jord, Freya, Baldr, and Sif were chosen to maximize the number of tips that match the species designation while minimizing those that do not. This algorithm was employed to account for the placement of some species outside their monophyletic groups, either due to incorrect species designations or other anomalies. Joint reconstruction was used to determine the number of ESP-containing orthogroups at each node, and these counts were then visualized against the total number of identified ESP-containing orthologs using the pheatmap function in R (41).

## Results

### Simulation Analysis

Across all conditions in our simulations, performance improves with genome count; larger datasets exhibit consistently higher precision at a given recall (**Figure 1**). When the number of genomes is small, precision declines rapidly as recall increases, indicating reduced robustness to incomplete observations in low-data scenarios. In contrast, datasets with several hundred genomes maintain high precision across most of the recall range, with degradation occurring primarily near perfect recall. Within each genome-count, variation across completeness profiles yields only modest differences in the precision–recall curves. Overall, the model performs strongly on idealized datasets.

**Fig. 1.**
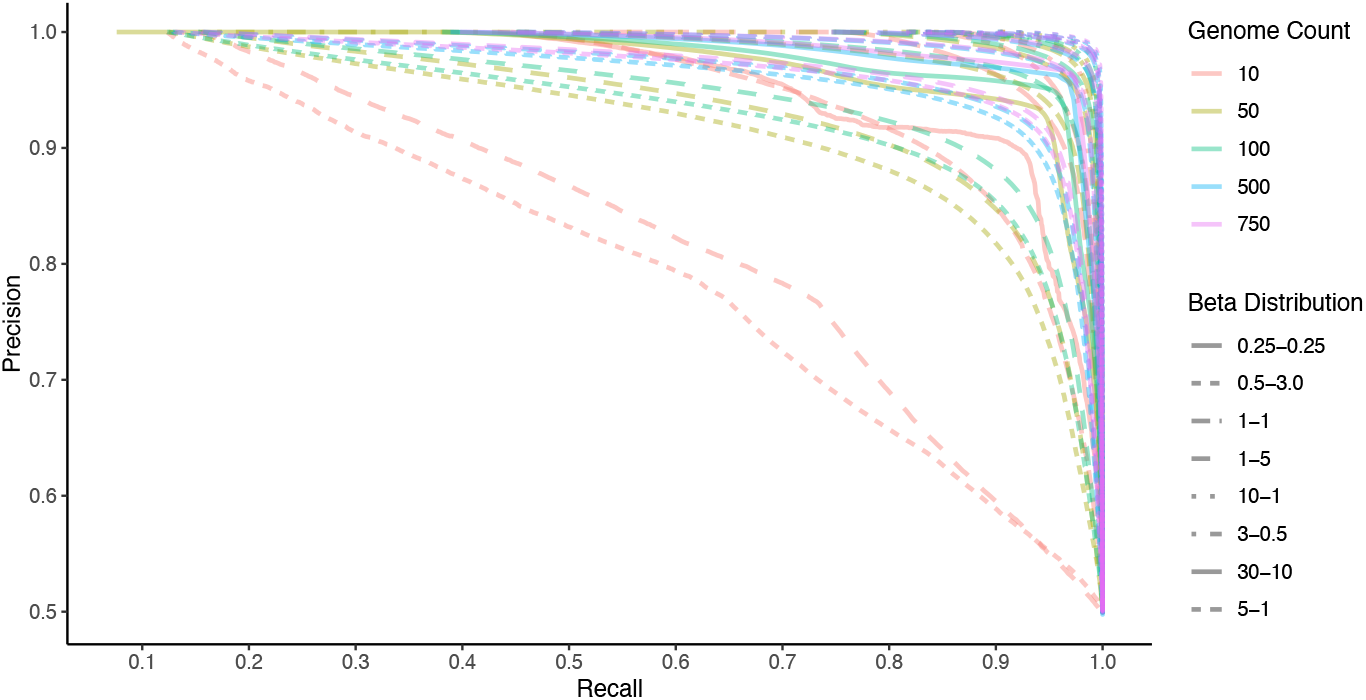
Precision and recall of presence/absence data simulated and run under our model at varying completeness profiles (Beta distributions) and number of genomes in the dataset. These metrics were only calculated on unobserved orthogroups; known presences do not contribute toward the precision and recall. This reflects the model’s performance in single-gene presence/absence imputation under idealized conditions.

### Empirical Evaluations

At the per-gene inference level, our model maintains favorable precision and recall across a wide range of completeness profiles (**Figure 3**). Although the model generally achieves higher precision and recall on incomplete datasets, these datasets contain more true positives available for detection. Under most completeness profiles, around 90% precision is obtainable at 40% recall. Overall, performance is generally lower than that of simulation analyses using a well-specified model. Still, it shows that reasonable per-gene inferences can be obtained when both the phylogeny and orthologous group assignments are inferred from real data.

Across both *α*- and *γ*-proteobacterial subclades (**Figure 2**), our model’s joint reconstruction and marginal probability thresholding consistently achieved higher recall for core genome inference than empirical thresholding approaches, while maintaining competitive precision. For the strict (present in 100%) definition of the core genome, the joint reconstruction exhibited near-perfect recall across most subclades, while the other methods frequently failed to recall anything (**Figure 4a, 4b**). Similar patterns were observed with the relaxed core definition (90% present) (**Figure 5a, 5b**); all methods showed modest improvements in recall, with our model achieving significantly higher precision.

**Fig. 2.**
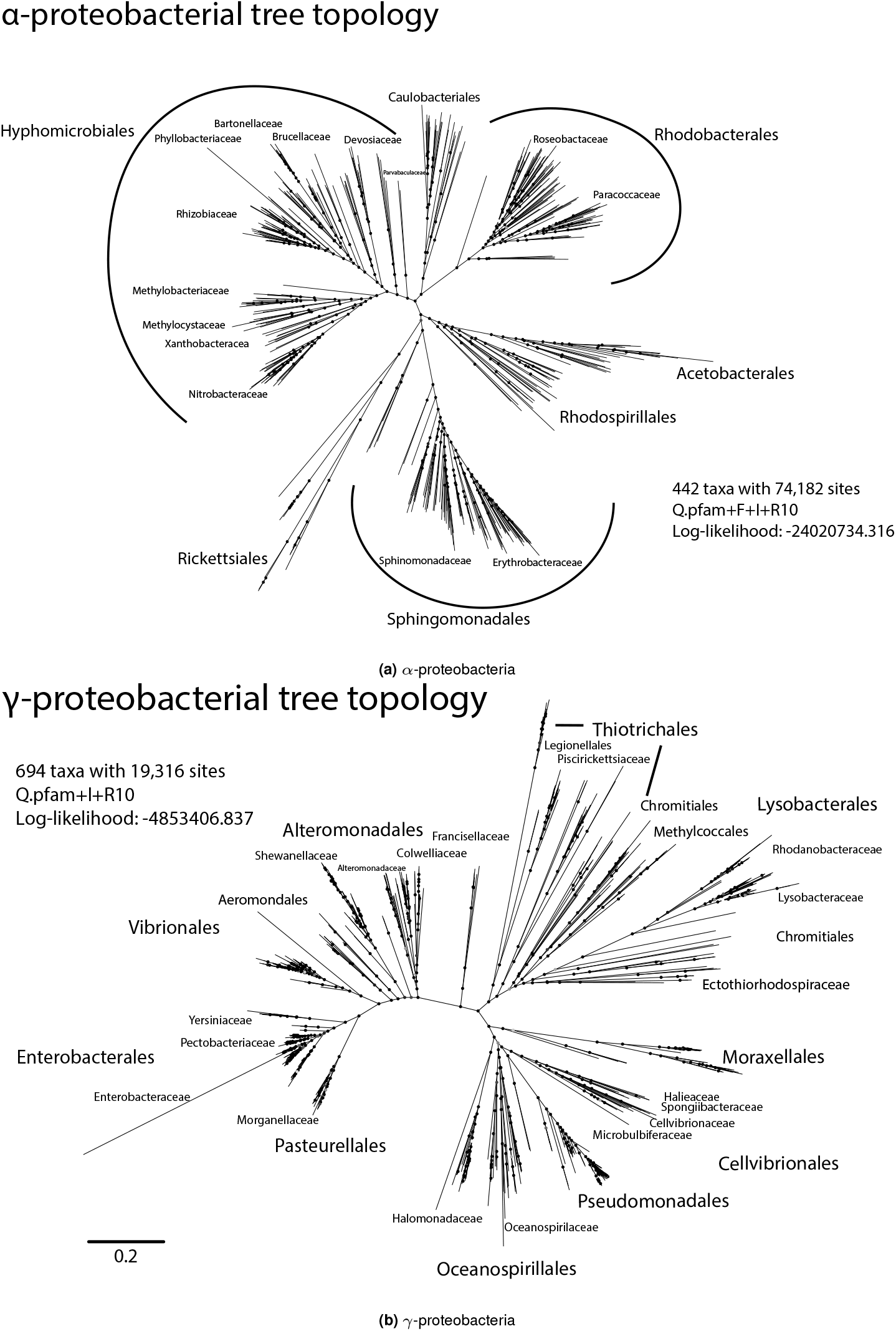
Phylogenetic relationships of *α*- and *γ*-proteobacteria. Maximum-likelihood phylogenies were reconstructed using concatenated single-copy genes present in 98% of taxa analyzed, and models were inferred using IQTree ModelFinder. Top: *α*-proteobacterial phylogeny inferred from 442 taxa and 74,182 aligned amino acid positions (log-likelihood = 2,402,0734.316). Major orders are indicated, including Hyphomicrobiales, Rhodobacterales, Caulobacterales, Rhodospirillales, Sphingomonadales, Rickettsiales, and Acetobacterales. Bottom: *γ*-proteobacterial phylogeny inferred from 694 taxa and 19,316 aligned amino acid positions (log-likelihood = 4,853,406.837). Major orders are labeled, including Enterobacterales, Vibrionales, Alteromonadales, Pseudomonadales, Oceanospirillales, Moraxellales, Cellvibrionales, Chromatiales, and Thiotrichales. Branch lengths represent the expected number of substitutions per site (scale bar shown). Trees are displayed as unrooted radial topologies for visualization of overall clade structure.

**Fig. 3.**
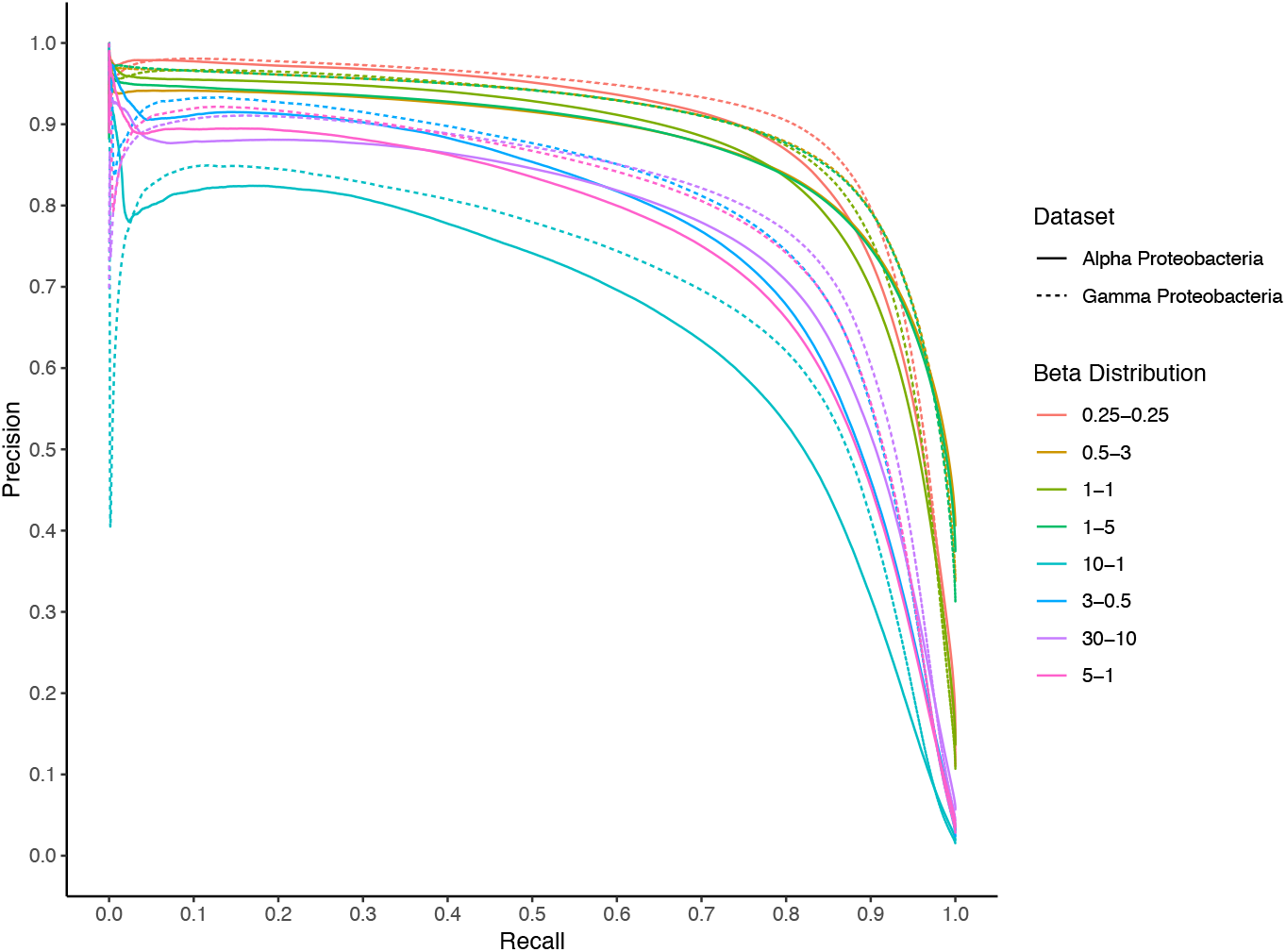
Precision and recall of our model for artificially depleted *α*-proteobacteria and *γ*-proteobacteria genomes under varying completeness profiles (Beta distributions). These metrics were only calculated on unobserved orthogroups; known presences do not contribute toward the precision and recall. This reflects the model’s performance in single-gene presence/absence imputation on two real datasets with some gene content masked.

**Fig. 4.**
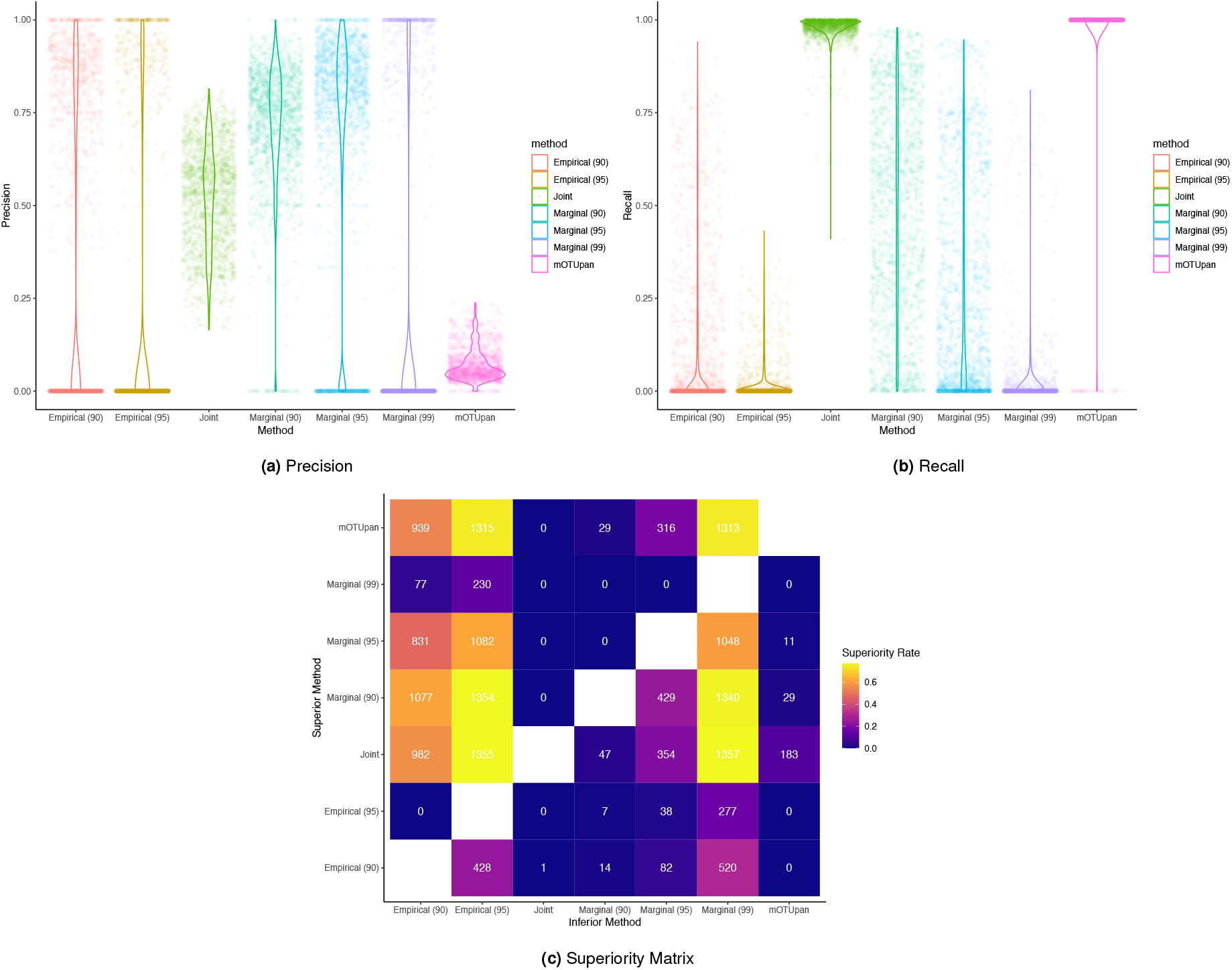
Performance metrics for strict core genome inference in the *α*- and *γ*-proteobacteria analyses. Core genomes were inferred for clades within the proteobacteria trees and predicted using: empirical thresholding (denoted “Empirical (*·*)”) of observed orthogroup presence/absence (i.e., the orthogroup is observed in 90% of incomplete genomes); joint maximum *a posteriori* reconstruction (denoted “Joint”) from our model (i.e., the maximum *a posteriori* reconstruction assigns the gene to all genomes in the clade); thresholding of marginal presence/absence probabilities (denoted “Marginal (*·*)”) from our model (i.e., the gene has greater than 90% marginal probability of presence in every genome in the clade); and mOTUpan core genome inference. Precision and recall for each method are shown in (a) and (b). The superiority matrix (c) summarizes pairwise comparisons, indicating the frequency with which each method is strictly better than the others (better in precision and recall).

**Fig. 5.**
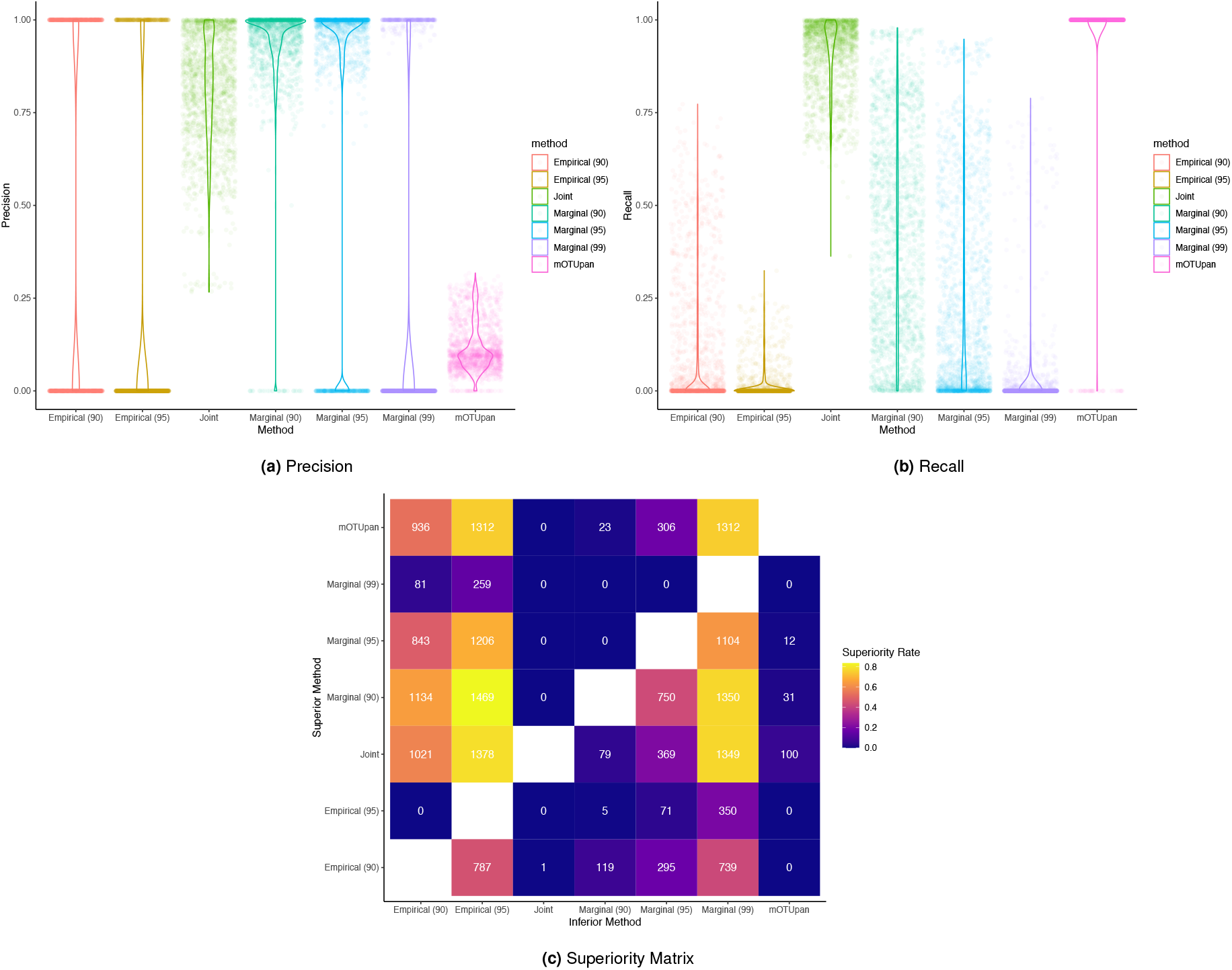
Performance metrics for relaxed core genome inference in the *α*- and *γ*-proteobacteria analyses. In these analyses, the definition of a “core” gene is any gene that is truly present in over 90% of the clade. Core genomes were inferred for clades within the proteobacteria trees and predicted using: empirical thresholding (denoted “Empirical (*·*)”) of observed orthogroup presence/absence (i.e., the orthogroup is observed in 90% of incomplete genomes); joint maximum *a posteriori* reconstruction (denoted “Joint”) from our model (i.e., the maximum *a posteriori* reconstruction assigns the gene to all genomes in the clade); thresholding of marginal presence/absence probabilities (denoted “Marginal (*·*)”) from our model (i.e., the gene has greater than 90% marginal probability of presence in every genome in the clade); and mOTUpan core genome inference. Precision and recall for each method are shown in (a) and (b). The superiority matrix (c) summarizes pairwise comparisons, indicating the frequency with which each method is strictly better than the others (better in precision and recall).

We used pairwise superiority matrices to simplify performance comparisons among methods (**Figure 4c, 5c**). Rather than assessing the benefits of greater/lesser recall at the cost of greater/lesser precision, we count the number of times each model is strictly superior to another (i.e., better in terms of both precision and recall). Across both strict and relaxed core definitions, the joint reconstruction output outperforms empirical thresholding and mOTUpan in the majority of comparisons, with marginal probability thresholding outperforming empirical thresholding, albeit with lower recall than mO-TUpan and joint reconstruction. Precision distributions show that while mOTUpan achieves high recall, it does so at the cost of substantially reduced precision, whereas our methods allow for higher precision.

### Asgardarchaea Analysis

To apply our model’s ancestral reconstruction capabilities, we used 741 Asgardarchaeal metagenomes of varying degrees of completeness. Two genomes were excluded from this analysis due to extremely poor gene content (more than 95% gaps in the alignment used to infer phylogeny). An outgroup of 206 TACK archaea was used to root the Asgard tree. The inferred phylogeny was monophyletic within TACK, allowing us to identify a potential root for ancestral state reconstruction (**Figure 6**). Eukaryotic signature proteins (ESPs) (39) were matched to eggnog annotations via BLAST matches, and the presence or absence of these specific orthologs across the Asgardarchaeal tree was observed (**Figure 6**). The ancestral reconstruction suggests significant gains and losses of ESPs after the divergence of the major Asgard groups, and often near the tips of the tree (**Figure 7c**). This is notably true in Heimdalarchaea and Hodarchaea, which have approximately the same number of ESPs in their common ancestor (158 ESPs in the reconstructed ancestor of Heimdalarchaea (**Figure 7a**), 166 in the reconstructed ancestor of Lokiarchaea (**Figure 7b**), and 169 in the reconstructed ancestor of Hodarchaea) as other major Asgard groups, but gain significantly more towards the tips (the maximum number of ESPs in any Asgard tip species after modeling is Heimdallarchaeota GCA011364945.1 with 241 ESPs). Based on this reconstruction, no Asgard group is able to claim more than 40% of identified Asgard ESPs in the common ancestor.

**Fig. 6.**
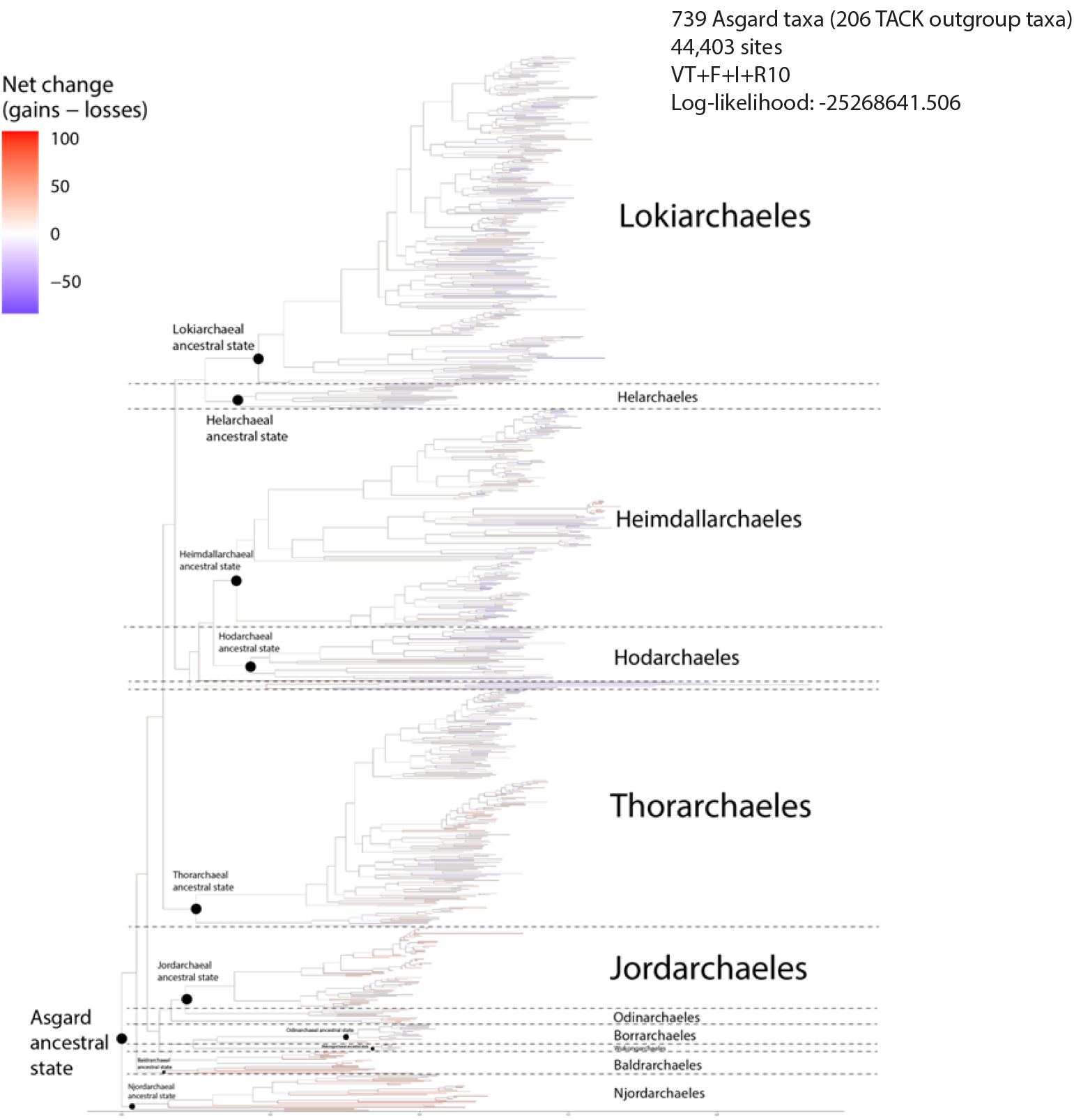
Asgard ancestral reconstruction of eukaryotic-specific protein gain-loss. Maximum-likelihood phylogeny of 739 Asgard taxa (including 206 TACK outgroup taxa not displayed in this tree) inferred from 44,403 aligned amino acid positions under the VT+F+I+R10 substitution model (log-likelihood = 25,268,641.506). Major Asgard clades are indicated, including Lokiarchaeales, Heimdallarchaeales, Hodarchaeales, Thorarchaeales, and Jordarchaeales, with additional subclades labeled where relevant. Net changes in eukaryotic-specific protein families were constructed using ancestral state inference and plotted using branch colors to indicate the direction and magnitude of the change in ESP presence. Black circles denote inferred ancestral nodes for major clades analyzed in Figure 7, including the overall Asgard ancestral state and order-level ancestors. Branch lengths are proportional to the expected number of substitutions per site.

**Fig. 7.**
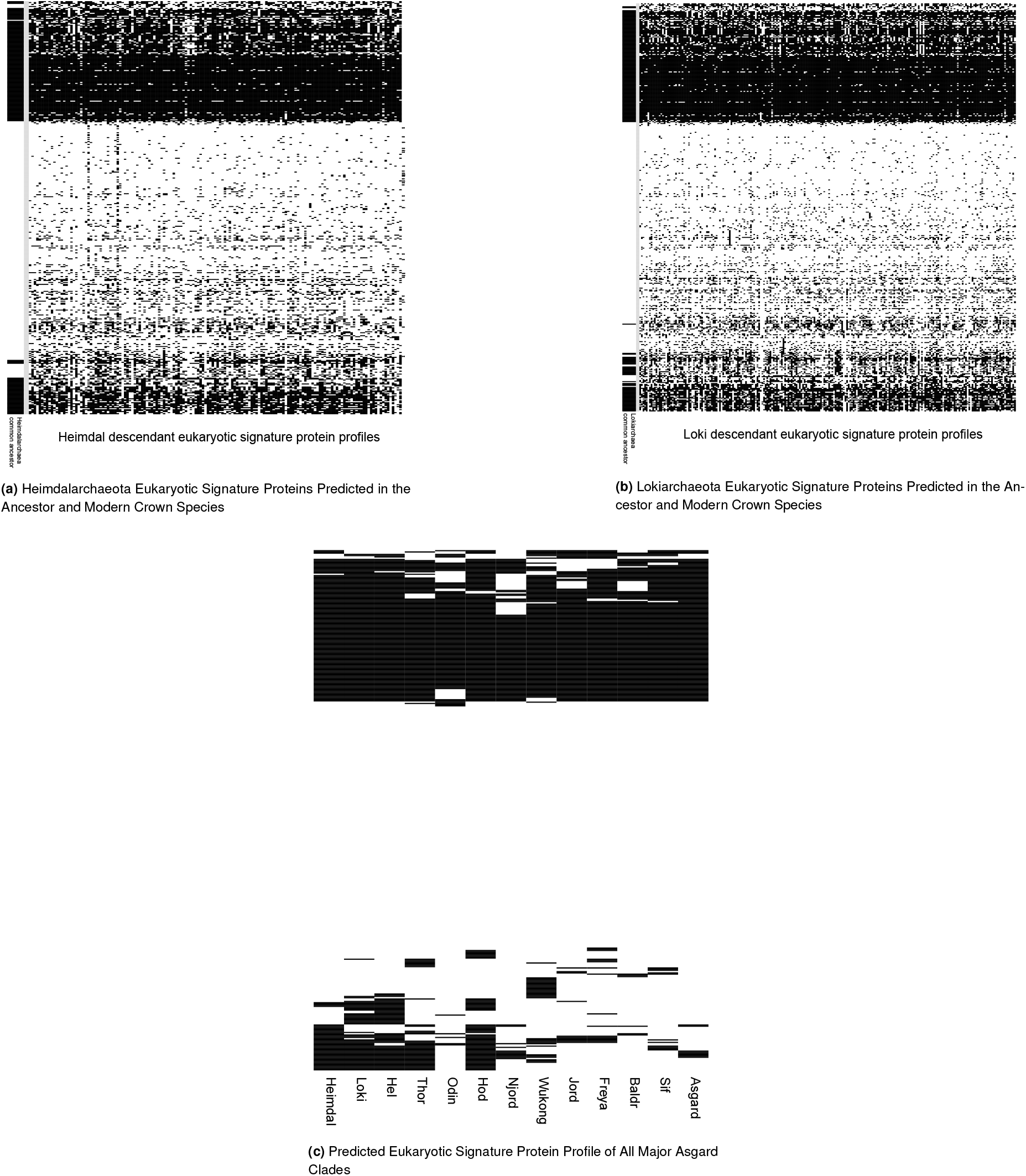
Ancestor and Crown Eukaryotic Signature Protein Profiles in Asgard Archaea after Occupancy Model Predictions of Crown gene content. Heatmaps summarize the presence–absence patterns of predicted eukaryotic signature proteins (ESPs) inferred using occupancy-based ancestral reconstruction of crown gene content. Black cells indicate predicted presence, white cells indicate absence. Rows correspond to individual ESP families; columns correspond to taxa or reconstructed ancestral nodes, as indicated below each panel. (a) ESP profiles for Heimdallarchaeota, showing predicted content in the Heimdallarchaeal ancestor alongside extant crown species. (b) ESP profiles for Lokiarchaeota, showing predicted content in the Lokiarchaeal ancestor compared to modern crown representatives. (c) Condensed ESP profile matrix summarizing reconstructed ancestral and crown gene complements across all major Asgard clades. Notably, in all Asgard ancestral cases a large proportion of ESPs are absent from the inferred ancestor despite their presence in one or more crown species members.

## Discussion

### Model Performance

Our initial simulation analyses indicate that the model performs very well when well-specified. Furthermore, its performance tends to improve as the number of taxa in the dataset increases. This behavior can be understood as arising from the model’s increasing number of imperfect “views” of each genome; each genome serves as an evolutionary replicate of the others. However, these data were simulated under the model itself, which is not the generative process that operates in real biological systems. Therefore, we sought to evaluate our model on high-quality biological data, for which we could be as confident as possible that gene presence/absence in each genome reflected true biological presence or absence.

We pulled from NCBI’s database of *α*- and *γ*-proteobacterial genomes, requiring that genomes be complete (circular, single-chromosome), RefSeq-standard, and fully annotated. Phylogenies were constructed from concatenated orthologs that had fewer than two copies across all genomes and were present in at least 90% of the included taxa. This procedure produced phylogenies for both *α*- and *γ*-proteobacteria (**Figure 2**), which were then used to evaluate model performance. We simulated incomplete data by artificially depleting the presence/absence matrices. A variety of completeness profiles were considered; although incompleteness arising from sequencing and assembly is likely more complex, this procedure preserves the generative process of evolutionary gene presence/absence followed by bioinformatic clustering.

At the per-gene level, our model achieves favorable precision and recall across a range of completeness profiles. The model generally performed better on *γ*-proteobacterial than on *α*-proteobacterial datasets (**Figure 1**). It is unclear whether this difference is driven by the extreme genomic reduction observed in some *α*-proteobacterial clades (42) or by the fact that the *γ*-proteobacterial dataset contains 252 more genomes. In general, precision increases as datasets become more incomplete; however, this effect is largely attributable to class imbalance rather than to the model itself. Nearly complete datasets contain fewer true positives that can be recovered. Nonetheless, across many completeness profiles, we can achieve approximately 40% recall at 90% precision, indicating that a substantial fraction of missing data can be recovered with reasonable confidence. In large-scale genomic inventory analyses, reporting these marginal probabilities may allow researchers to assign confidence to gene presence in individual genomes.

For core genome inference, our approach substantially outperforms alternative methods (**Figure 4, 5**). Joint reconstruction more reliably recovers core genomes than competing approaches, albeit with lower precision, whereas marginal-probability-based thresholding achieves superior precision and recall relative to empirical thresholding based on observed presence/absence. Although our methods outperform the others we considered here, reconstructing the core genome from incomplete data remains a very difficult problem. For example, four taxa in our *γ*-proteobacterial dataset are missing COG0858: *Ahniella affigens, Billgrantia tianxiuensis, Luteimonas granuli*, and *Pseudomonas purpurea*. These taxa are not closely related on the phylogenetic tree, and in all four genomes, the gene has been pseudogenized through one or more point mutations. This orthogroup is typically present as a single copy and encodes the protein rbfA, which assists in maturation of the 30S ribosomal subunit in *Escherichia coli* (43). A reasonable prediction by our algorithm (or even by a biologist) would be that this gene is present in all sampled *γ*-proteobacterial genomes, but it is apparently absent in any functional form. Further modifications to our model may improve performance, but recent evolutionary events of this kind will likely remain difficult to predict. In its strictest formulation, core genome inference and gene presence/absence estimation are inherently difficult; some irreducible error is likely unavoidable in the strict core genome inference problem.

We would like to emphasize that although mOTUpan performs poorly in our evaluations, the deep-time phylogenetic relationships considered here are substantially different from the use case for which it was designed. Indeed, its original benchmarks focused on within-genus datasets of *Prochlorococcus* genomes (18). In such settings, genome-content heterogeneity may be limited enough to justify their modeling decisions. We anticipate that our model would also perform well in this setting.

### Asgardarchaea Analysis

Eukaryotes are thought to root within the Asgardarchaea, more specifically as a sister to either the Heimdalarchaea or Hodarchaea, and the divergence of these species occurred approximately 2 billion years ago (44, 45). Previous analyses (46, 47) that examined the ancestral state of Asgardarchaea focused primarily on the metabolic nature of Heimdal- and Hodarchaea, rather than the proteins in their genomes that were previously thought to be eukaryote-specific (ESPs). These analyses suggest that the ancestors of Heimdal and Hod had metabolic capabilities similar to those of the crown species in those clades, consistent with our ancestral state reconstructions of those metabolic pathways. However, the metabolic conservation between crown and ancestral Asgard metabolic capabilities was not carried through to the eukaryotic-specific proteins found in crown Asgards (**Figure 7c**). These proteins were previously considered present exclusively in eukaryotes, until their identification in at least one Asgardarchaeal species, and are thought to have been inherited by eukaryotes from an Asgard ancestor. Our reconstruction indicates that all Asgard common ancestors had a similar number of ESPs (40% of the total ESPs identified in Asgard genomes), and although Hodarchaea have the most ESPs, there remains a significant gap between their ancestral and modern gene content. As shown in (**Figures 7a,7b**), many of these ESPs are present in modern crown Asgardarchaea but are absent in ancestral-state reconstructions, suggesting patchy gain and loss of these proteins across all Asgardarchaea families. Notably, functional roles of gene families in the ancestral-state reconstructions included actin regulation, ESCRT transport, and cell cycle checkpoint proteins, suggesting that ancestral Asgardarchaea were capable of membrane remodeling similar to that observed in *Lokiarchaeum ossiferum*. An analysis of gene families specific to the ancestor of Heimdalarchaea that were absent from the ancestor of all Asgard identified 33 ESPs with diverse functions, including vesicular transport, GTPase, and gyrase protein families. Although some discrepancies may be attributable to errors in the phylogeny, this pattern is observed throughout the tree. Further refinement of the model, as well as more complete genomes of diverse Asgard, will be able to improve this analysis to create a more accurate picture of the shared ancestor between Asgardarchaea and eukaryotes, though it is worth noting that Asgardarchaea are not fixed fossils in time, and it makes sense that they would continue to evolve with respect to gene content just as eukaryotes did.

### Future Directions

In this work, our model design was guided primarily by interpreting the phylogeny as a belief network rather than by strict adherence to standard phylogenetic formulations. Our departures from classical phylogenetic models are relatively modest, most notably our decision not to use the stationary distribution of the transition matrix ***Q*** as the root distribution and our choice of transition probabilities that reflects agnosticism in the limit of time, rather than encoding dynamics. We anticipate that further deviations from classical phylogenetic assumptions may yield additional improvements in model performance. More broadly, we believe phylogenetic occupancy models represent a promising approach to improving the inference of gene presence-absence in incomplete genomic datasets. In addition to performing better on core genome inference for deep-time evolutionary datasets, these models also enable inference of the presence of individual genes in particular genomes and ancestral state reconstruction under data uncertainty. Although our phylogenetic occupancy model improves on existing approaches, several plausible directions could yield further improvements. These include:

- Explicitly accounting for uncertainty in phylogenetic structure. Reasonable approaches include “cut” Bayesian approaches and fully Bayesian approaches. Cut Bayesian models use the posterior distribution from an initial analysis as the true posterior distribution for a subsequent analysis (48). These methods are particularly useful when the secondary analysis contains information you do not want to flow into the posterior distribution of the first analysis. This is reasonable here because we may not believe that gene presence-absence should strongly sway our beliefs about the evolutionary relationships between organisms. Alternatively, fully Bayesian approaches may be reasonable and have been applied to several mixed-data scenarios (49).
- Incorporating gene co-occurrence information into the model. This poses a serious challenge, as it requires defining conditional distributions over binary strings rather than simple binary states. In the simulations we explored, this would involve defining conditional distributions over 2^5000^ potential states. Approaches for phylogenetic modeling in this context do exist (50, 51), but remain computationally difficult. A potentially interesting approach would be to infer small cliques of genes that influence one another’s presence/absence, which could help prevent the state space from becoming prohibitively large.

## Code Availability

All the code required to run the phylogenetic occupancy model can be found at https://github.com/Wesley-DeMontigny/Phylogenetic-Occupancy-Model. Additional files can be found at https://doi.org/10.5281/zenodo.18786143.

## ACKNOWLEDGEMENTS

This research is funded in part by the Gordon and Betty Moore Foundation through Grant GBMF11481 (https://doi.org/10.37807/GBMF11481) to the University of Maryland (UMD). WCD was partially funded through a TAship from UMD. We would like to thank the attendees of the *Archaea: Ecology, Metabolism and Molecular Biology* Gordon conference for the helpful discussion on the model.

## Supplemental Materials: Core Genome Inference with Phylogenetic Occupancy Modeling

**Fig. S1.**
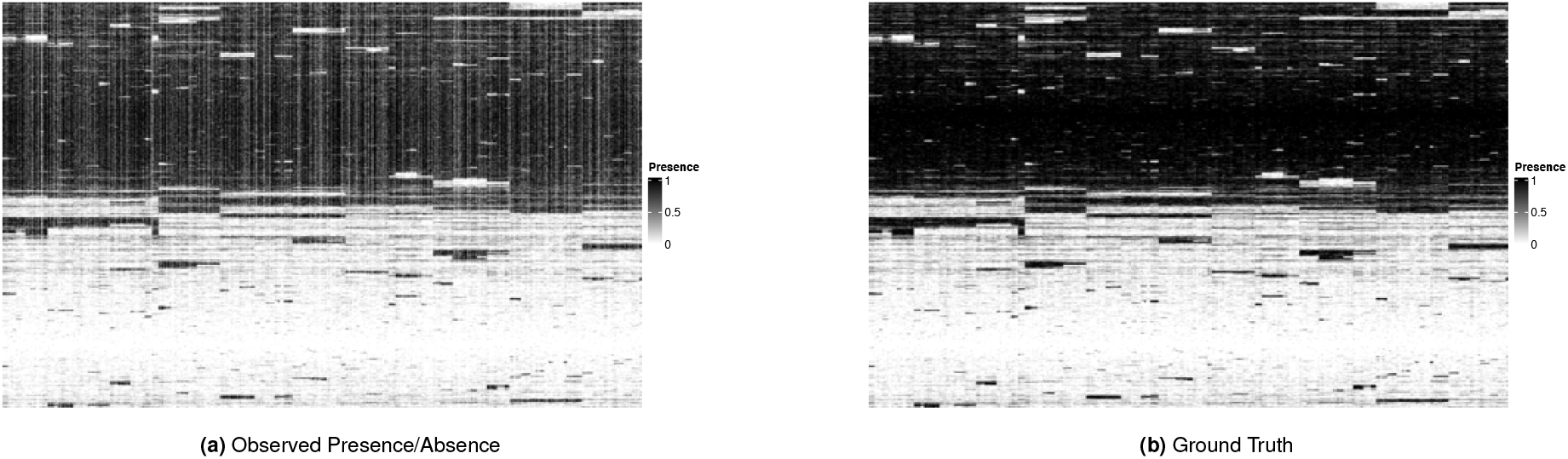
Example of observed data (a) and ground truth (b) from our simulation analyses. Specifically, this simulation had 750 genomes and 5,000 genes and was depleted according to a Beta(5, 1) distribution.

**Fig. S2.**
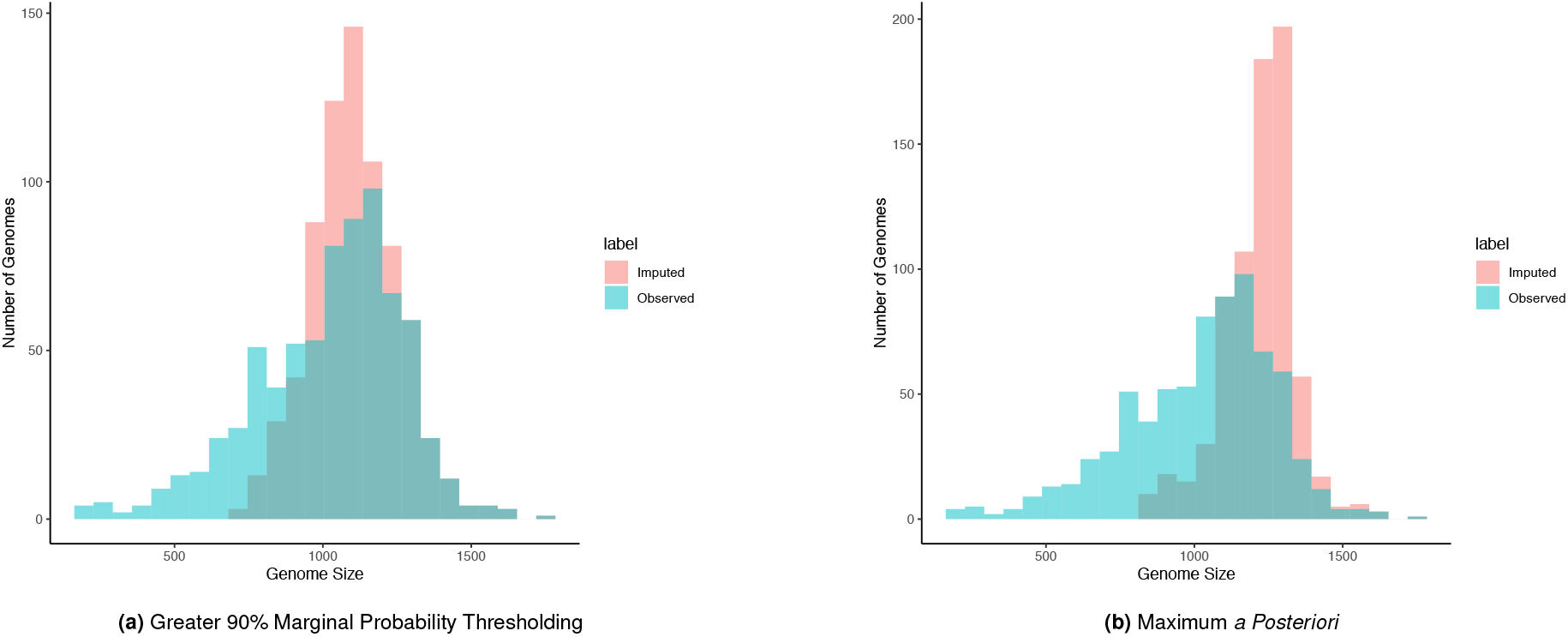
Histogram representing frequencies of genome size (in orthogroup counts) of Asgardarchaea genomes before and after gene imputation using (a) 90% marginal probability thresholding and (b) maximum *a posteriori* reconstruction.

